# 3D bioprinting of high cell-density heterogeneous tissue models through spheroid fusion within self-healing hydrogels

**DOI:** 10.1101/2020.05.21.103127

**Authors:** Andrew C. Daly, Matthew D. Davidson, Jason A. Burdick

**Affiliations:** Department of Bioengineering, University of Pennsylvania, Philadelphia, PA, USA

## Abstract

Cellular models are needed to study human development and disease *in vitro*, including the screening of drugs for toxicity and efficacy. However, current approaches are limited in the engineering of functional tissue models with requisite cell densities and heterogeneity to appropriately model cell and tissue behaviors. Here, we develop a new bioprinting approach to transfer spheroids into self-healing support hydrogels at high resolution, which enables their patterning and fusion into high-cell density microtissues of prescribed spatial organization. As an example application, we bioprint induced pluripotent stem cell-derived cardiac microtissue models with spatially controlled cardiomyocyte and fibroblast cell ratios to replicate the structural and functional features of scarred cardiac tissue that arise following myocardial infarction, including reduced contractility and irregular electrical activity. The bioprinted *in vitro* model is combined with functional readouts to probe how various pro-regenerative microRNA treatment regimes influence tissue regeneration and recovery of function as a result of cardiomyocyte proliferation. This method is useful for a range of biomedical applications, including the development of precision models to mimic diseases and for the screening of drugs, particularly where high cell densities and heterogeneity are important.

## Introduction

Cells possess the remarkable capacity to self-organize into multicellular spheroids *in vitro*, including from stem cells and into specialized organoid structures ^*1*^. These spheroids are being used to emulate organs such as the intestine ^2^, liver ^3^, kidney ^4^, brain ^5^, and heart ^6^ to study human development and disease *in vitro*. In these formats they are promising as drug screening platforms due to their superior predictability and physiological structure and function when compared to traditional monolayer cultures ^7-10^. Indeed, the organotypic cell densities in spheroids enhances the cell-cell and cell-extracellular matrix (ECM) interactions that are required to maintain cellular differentiation and phenotype, whereas these interactions are limited in traditional 2D cultures^11-14^. High cell-densities are necessary to accurately recapitulate many pathological disease states such as cancer or fibrosis where perturbed cell-cell and cell-ECM interactions are central to disease progression. For example, spheroid cultures have enabled the engineering of heart, liver, and lung fibrosis models ^6, 15-18^, and the presence of oxygen gradients with high cell densities mimics the cancer microenvironment ^19-21^.

While spheroid cultures hold tremendous potential due to their organotypic cellular densities, it is difficult to control their patterning across larger length scales, to introduce the heterogeneity and resulting function of many tissues. This limits their utility as *in vitro* models where patterns of cells and ECM are key aspects of development or the progression of disease. For example, during fibrosis of the heart, liver and lung ^22, 23^, non-functional fibrotic tissue persists within healthy parenchyma, promoting further scarring and eventual organ failure. The development of biofabrication tools to control spheroid assembly across larger length scales through directed placement and fusion could improve their physiological relevance for a range of biomedical applications.

Biofabrication technologies, such as bioprinting, enable programmed assembly of cells into complex 3D geometries ^24, 25^. However, these technologies have been predominantly designed to process cells embedded within hydrogels, which limits cell-cell interactions and results in low cell density constructs. This has motivated the development of biofabrication technologies tailored specifically for spheroids, often termed bioassembly, which leverages the capacity of spheroids to fuse through liquid-like coalescence to minimize adhesive-free energy via cellular remodeling of the interface ^26-28^. In an early approach, extrusion bioprinting was adopted to allow spheroids to fuse together into larger tissues strands, which could then be extruded through a microcapillary ^29^. More recent bioassembly advances enable the direct processing of single spheroids, offering improved control over spheroid patterning and overall geometries. For example, spheroids can be aspirated and then skewered onto supporting metallic needles (i.e., Kenzan method) for fusion ^30^. Hydrogels have also been used to support aspiration based methods, through the sequential layering of an uncrosslinked hydrogel precursor and spheroids, followed by layer crosslinking ^31^. Despite advances with these bioassembly technologies, significant challenges remain, particularly to support the patterning of complex 3D structures without the need to disrupt spheroids mechanically or to use external stimuli to initiate hydrogel crosslinking. Thus, the development of improved and simplified platforms to pattern spheroids in 3D are needed.

Here, we report a new method where spheroids can be translated through a shear-thinning hydrogel that self-heals to receive and hold the spheroid in 3D space, including for the directed fusion between spheroids to form high-cell density microtissues with defined shapes that can then be removed from the hydrogel. The self-healing properties of the support hydrogel enable precise positioning of spheroids (up to 10% of spheroid diameter) and high spheroid viability post-printing (∼95%) while the viscoelastic and non-adhesive nature of the self-healing hydrogel facilitates controlled fusion between adjacent spheroids into prescribed and stable geometries during culture. To demonstrate the utility of this new bioprinting method, we develop a cardiac disease model that mimics post-myocardial infarction (MI) scarring, by bioprinting microtissues containing spatially controlled densities of induced pluripotent stem cell (iPSC)-derived cardiomyocytes and fibroblasts. The bioprinted model replicates cardiac pathologies that arise post-MI (reduced contractile output and electrical synchronization) and we use the model to probe miRNA therapeutics to improve cardiac regeneration and function as a result of cardiomyocyte proliferation.

## Results and Discussion

### 3D bioprinting spheroids within self-healing hydrogels

To enable the controlled patterning of spheroids, we developed a new method for the bioprinting of spheroids within self-healing support hydrogels, which involves: (i) vacuum aspiration of a spheroid in a media reservoir, (ii) direct transfer of the spheroid into a support hydrogel due to its shear-thinning properties, and (iii) deposition of the spheroid at any location in the hydrogel with vacuum removal and self-healing of the hydrogel (Fig. 1a i-iii, Supplementary Movie 1). This process is enabled by the separation of the media reservoir containing the spheroids from a second reservoir containing the support hydrogel by a narrow channel for spheroid transport between the two reservoirs (Supplementary Fig. 1a, Supplementary Movie 1). This approach avoids surface tension effects that can significantly reduce cell viability when transferring cells and spheroids between liquid and air interfaces^31^. The support hydrogel used is based on the assembly of hyaluronic acid (HA) modified with either adamantane (Ad) or β-cyclodextrin (CD) (Supplementary Fig. 2), resulting in shear-thinning and self-healing properties, as represented by a reduction in hydrogel viscosity with increasing shear rate and a recovery of storage modulus during cycles of high to low strain (Fig. 1b). This supramolecular hydrogel was developed previously for applications in the delivery of therapeutics to tissues and as a bioink for extrusion printing^32-34^.

**Figure 1:**
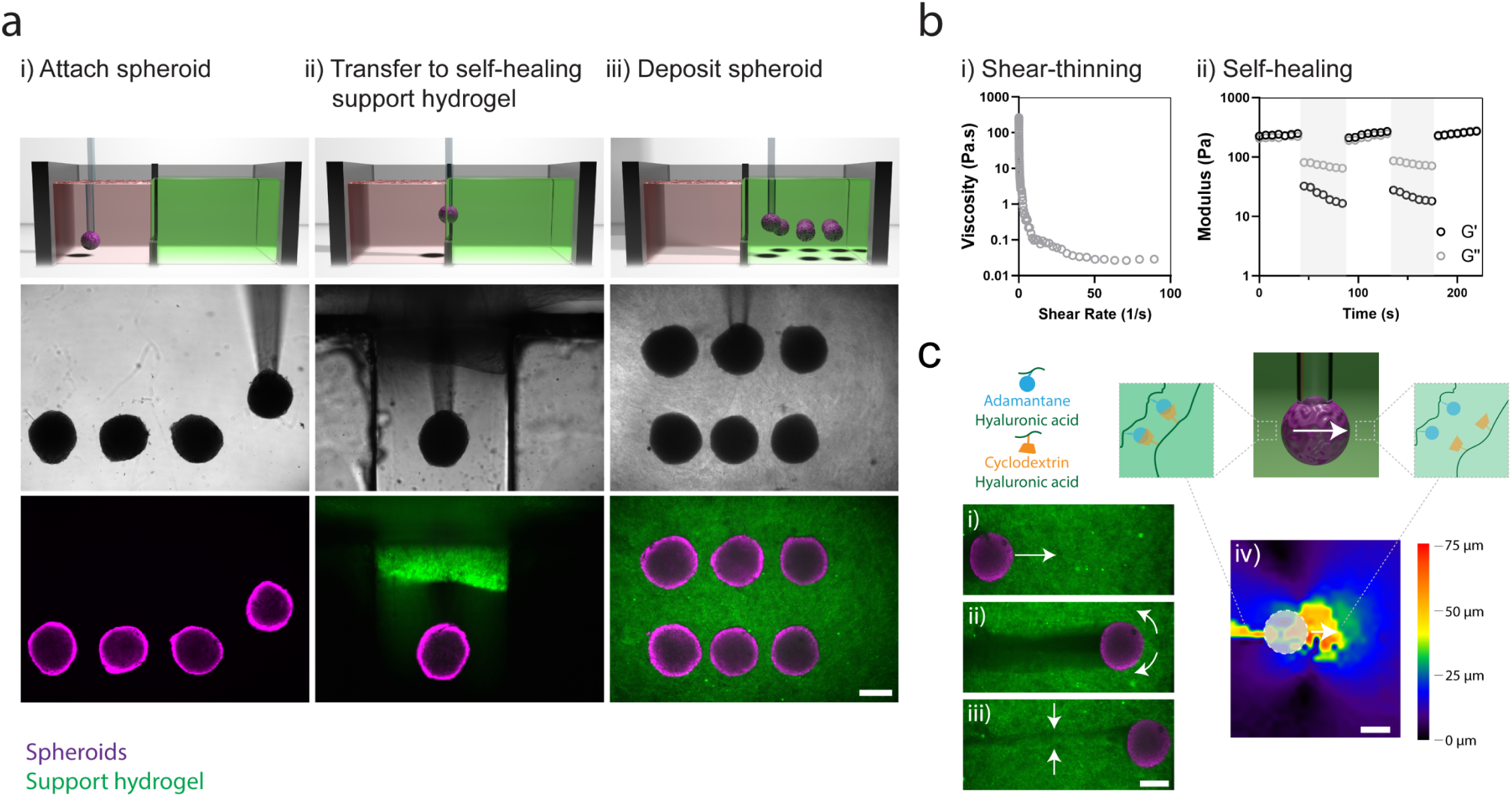
3D bioprinting spheroids in self-healing support hydrogels: **(a)** Schematic (top), brightfield images (middle), and fluorescent images (bottom) demonstrating, (i) MSC spheroid aspiration in a media reservoir, (ii) spheroid transfer into a self-healing support hydrogel (FITC-labeled), and (iii) spheroid deposition within the support hydrogel through removal of vacuum from the capillary tip. **(b)** Rheological characterization of a guest-host support hydrogel (3 wt%) demonstrating, (i) shear-thinning properties – decreased viscosity with continuously increasing shear rates (0–100 s^-1^) and (ii) self-healing properties – storage and loss modulus recovery (G’ & G’’) through low (0.5% strain, 10 Hz) and high (shaded, 100% strain, 10 Hz) strain cycles. **(c)** Reversible interactions between guest (adamantane, blue) and host (β-cyclodextrin, orange) modified hyaluronic acid of the support hydrogel (containing FITC microparticles) enable, (i-ii) local yielding of the support hydrogel under shear during spheroid translation, and (ii-iii) rapid healing of the support hydrogel after spheroid translation. (iv) Displacement mapping of the support hydrogel (spheroid noted as dashed circle) demonstrating local motion of the hydrogel in front of and behind the spheroid during spheroid translation. All scalebars 250 µm.

**Figure 2:**
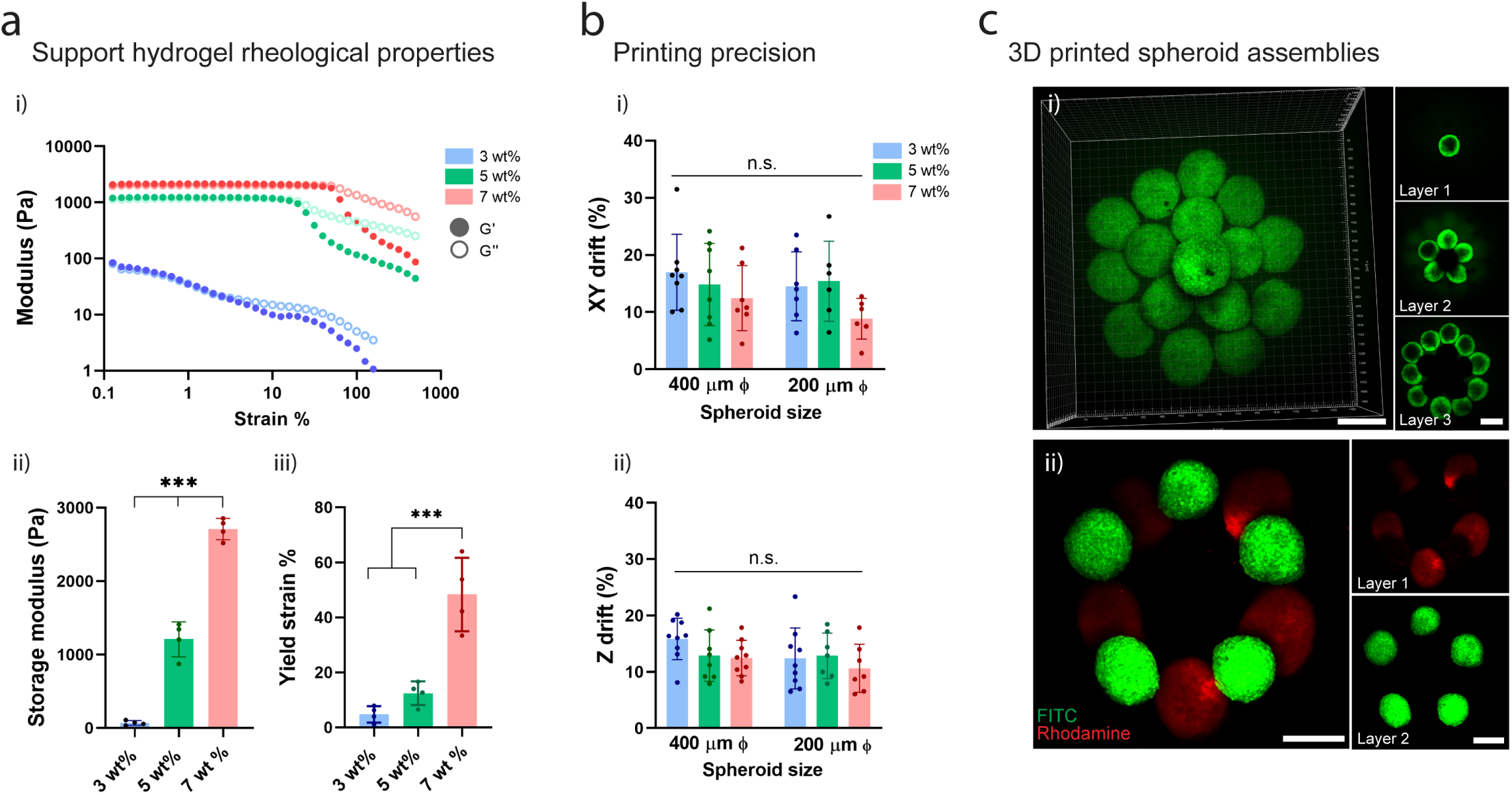
Support hydrogel rheological properties and 3D bioprinting precision. **(a)** (i) Shear-yielding in guest-host support hydrogels with increasing strain (0.1–500%, 10 Hz) at varied macromer concentrations (3, 5, 7 wt%) (storage modulus G’, closed circles – loss modulus G’’, open circles). (ii) Storage modulus at low strain (0.5%) at varied macromer concentrations (n=4, mean ± s.d, one-way ANOVA, *** p<0.001). (iii) Yield point (strain at G’/G’’ crossover) at varied macromer concentrations (n=4, mean ± s.d, one-way ANOVA, *** p<0.001). **(b)** (i) Bioprinting precision in the XY plane (XY drift %) for 200 and 400 µm diameter spheroids (n=6-8, mean ± s.d, one-way ANOVA, n.s. not significant). XY drift % = post-printing spheroid drift / spheroid diameter (see Supplementary Fig. 4a for schematic of measurement). (ii) Bioprinting precision in the Z plane (Z drift %) for 200 and 400µm diameter spheroids (n=7-9, mean ± s.d, one-way ANOVA, n.s. not significant). Z drift % = post-printing spheroid drift / spheroid diameter (see Supplementary Fig. 4b for schematic of measurement). **(c)** 3D bioprinted MSC spheroids within a support hydrogel (3 wt% macromer concentration) into either (i) a multi-layer cone shaped geometry (FITC-labeled spheroids) or (ii) layered rings of distinct MSC spheroid populations (FITC- or rhodamine-labeled) within the support hydrogel. All scalebars 250µm.

To visualize hydrogel motion during spheroid bioprinting, we embedded FITC-microparticles within the support hydrogel and used particle-image-velocimetry (PIV) analysis to track particle motion. Shear-thinning properties allowed the support hydrogel to be locally disrupted to accommodate spheroid translation (Fig. 1c i-ii, Supplementary Movie 2) and self-healing properties allowed the hydrogel to reassemble to hold the spheroid in place following the release of vacuum (Fig. 1c iii, Supplementary Movie 2). Quantitative mapping of relative particle motion indicated that the hydrogel yielded predominantly at the front of the translating spheroid with limited hydrogel movement further than ∼200 µm from the spheroid (Fig. 1c iv, Supplementary Fig. 3). Due to inherent hydrogel properties, the bioprinting process could be repeated, allowing the deposition of multiple spheroids within the hydrogel at any location in 3D space not previously occupied (Fig. 1aiii).

**Figure 3:**
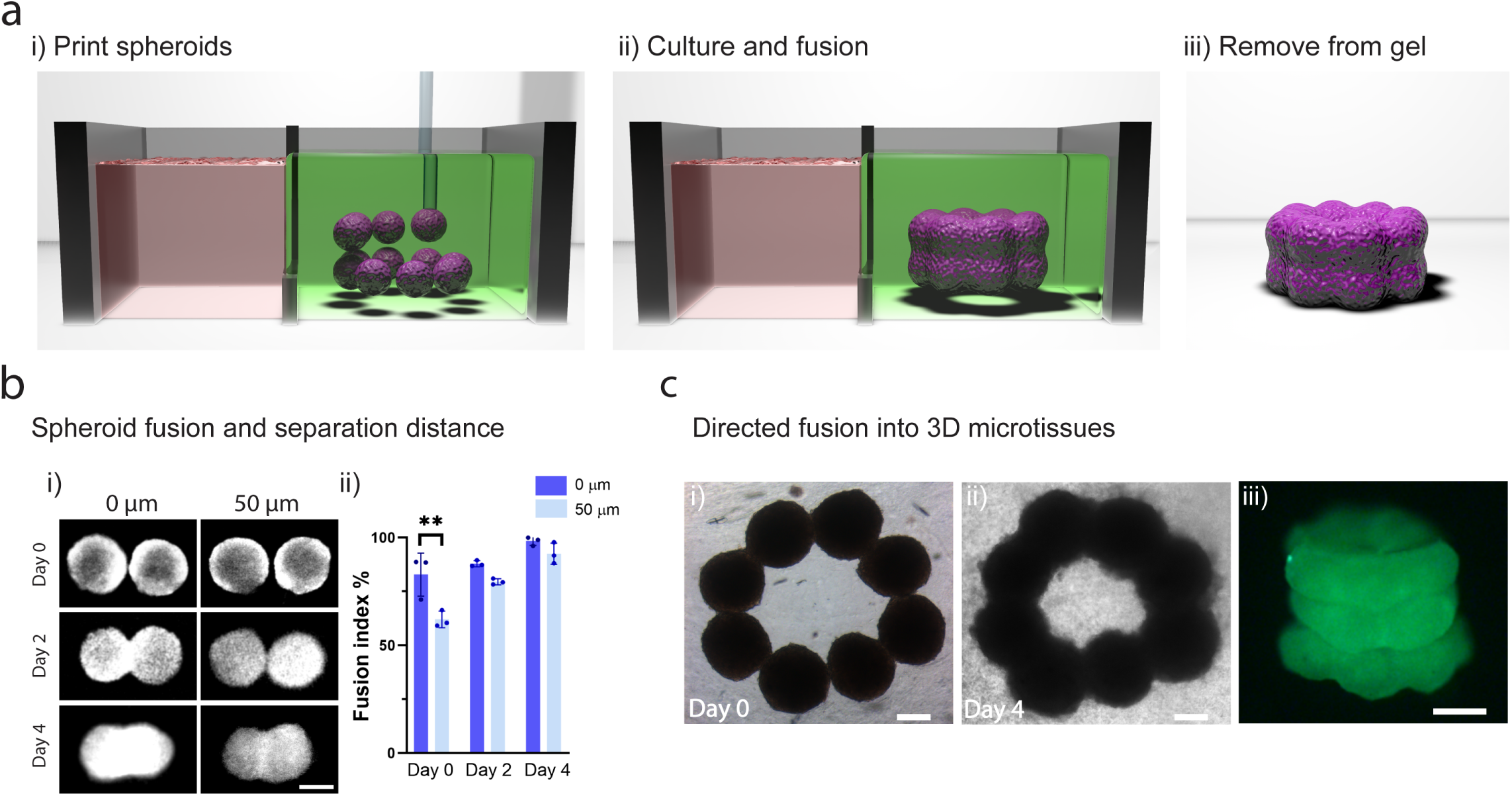
3D bioprinting microtissues through directed spheroid fusion. **(a)** (i) Schematic demonstrating the bioprinting of spheroids into rings within the support hydrogel, (ii) spheroid fusion and microtissue formation within the support hydrogel during culture, and (iii) removal of the microtissue via the reversibility of the support hydrogel crosslinking. **(b)** (i) Spheroid fusion over 4 days as a function of initial separation distance (0 or 50 µm). (ii) Fusion index (%) (see Supplementary Fig. 6 for schematic of measurement) over 4 days of culture (n=3, mean ± s.d, one-way ANOVA, **p<0.01 (Day 0; 0µm vs. 50µm p=0.0025)). **(c)** (i-ii) Brightfield images of spheroid fusion into microtissue rings over 4 days of culture. (iii) 3D bioprinted microtissue tube composed of 3 layers of fused spheroids (after 4 days of culture and then removal from the support hydrogel). All scalebars 200µm.

Having illustrated the ability to bioprint spheroids inside self-healing support hydrogels, we next explored how the hydrogel rheological properties influenced bioprinting precision. Increasing the polymer concentration from 3 to 7 wt% resulted in hydrogels with higher storage moduli that required higher strains to initiate yielding (Fig. 2a i-iii). To assess bioprinting precision, we developed assays where the XY and Z locations of a bioprinted spheroid is tracked during culture in the support hydrogel post-printing, which was performed for two spheroid sizes (200 and 400 µm Ø) and across polymer concentrations from 3 to 7 wt% (Supplementary Fig. 4). For XY precision, we measured drift distances of 8-15% of the spheroid diameter (or 15-60 µm) for all polymer concentrations (Fig. 2b i). For Z precision, we measured similar drift distances (10-15%) for all polymer concentration (Fig. 2b ii). These findings indicate high precision of bioprinted spheroids, supporting the approach for the high resolution bioprinting of spheroids into patterns.

**Figure 4:**
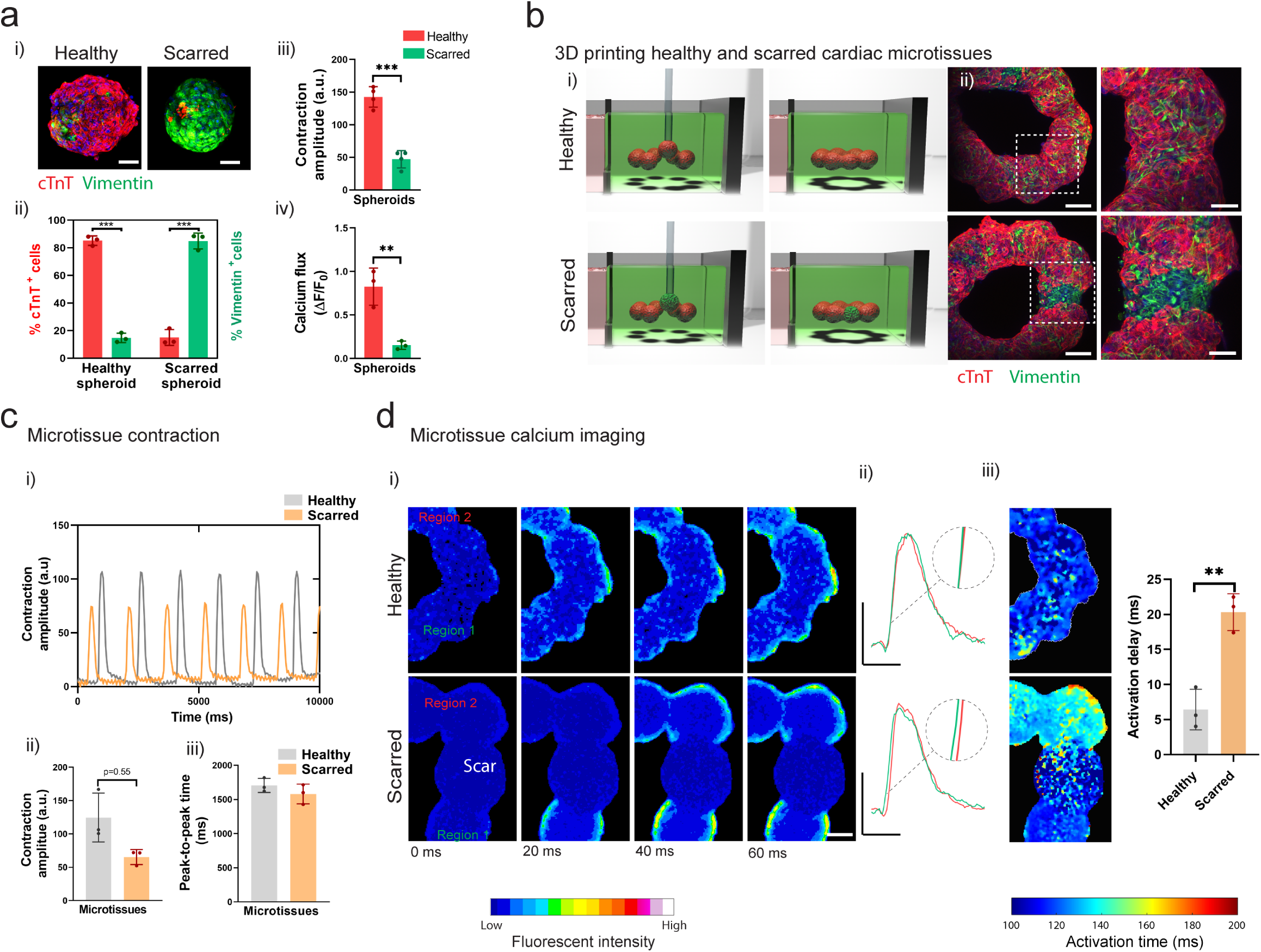
3D bioprinting cardiac microtissues for disease modeling applications. **(a)** (i) Development of healthy and scarred spheroids through mixing iPSC-derived cardiomyocytes (iPSC-CMs) with cardiac fibroblasts (CFs) at defined ratios of cell numbers (4:1 for healthy; 1:4 for scarred). Images taken 72 hours after cell seeding. Scalebar 50µm. (ii) Quantification of cellular composition through staining for cardiac troponin-T (cTnT) (iPSC-CMs) and vimentin (CFs) (n=3, mean ± s.d, two-sided student t-test, *** p<0.001, n=1 donor (A)). (iii) Contraction amplitude in healthy and scarred cardiac spheroids (n=4, mean ± s.d, two-sided student t-test, *** p<0.001, n=1 donor (A)). (iv) Normalized calcium flux amplitude (ΔF/F_o_) in healthy and scarred spheroids measured through fluorescent optical mapping (n=3, mean ± s.d, two-sided student t-test, ** p<0.01 (p=0.006), n=1 donor (A)). **(b)** (i) Schematic of the 3D bioprinting of healthy and scarred cardiac microtissue rings (healthy microtissues - 8 healthy spheroids; scarred microtissues - 7 healthy spheroids and 1 scarred spheroid) and (ii) Immunofluorescence staining for cTnT and vimentin in healthy and scarred cardiac microtissues after 5 days of fusion within the support hydrogel. Scalebar 100 µm (insets 50 µm). **(c)** (i) Contraction profiles of healthy and scarred cardiac microtissues following removal from the support hydrogel after 5 days of culture. (ii) Contraction amplitude (a.u. absolute units) at 5 days (n=3, mean ± s.d, two-sided student t-test, p=0.055, n=1 donor (A)). (iii) Peak-to-peak time (ms) at 5 days (n=3, mean ± s.d, two-sided student t-test n=1 donor (A)). **(d)** (i) Calcium mapping in healthy and scarred cardiac microtissues after 5 days of culture, where each image represents a 20 ms frame. Scalebar 100µm. (ii) Representative calcium traces from regions 1 and 2 in healthy and scarred cardiac microtissues, scale 250 ms (for scarred microtissues regions 1 and 2 are defined as spheroids directly adjacent to scarred region, for healthy microtissues regions 1 and 2 are defined as spheroids directly adjacent to a random healthy spheroid). Scalebars 0.5 ΔF/F_o_ (y),500 ms (x). (iii) Activation maps of healthy and scarred cardiac microtissues, and activation delay (ms) (difference in activation time (ms) between regions 1 and 2) in healthy and scarred cardiac microtissues, scalebar 100µm. Activation time is defined as the time taken for the calcium signal to reach 50% of its peak value during a single upstroke. (n=3, mean ± s.d, two-sided student t-test, ** p<0.01 (p=0.0035), n=1 donor (A)).

Next, we evaluated the viability of cells within bioprinted spheroids 24 hours post-printing for all polymer concentrations (Supplementary Fig. 5). At lower polymer concentrations (3 and 5 wt%) high cell viabilities (∼90-95%) were found, whereas a significant decrease in cell viability (∼87%) was observed at the highest polymer concentration (7 wt%). Lower polymer concentrations likely reduce the impact of shear forces during spheroid translation, resulting in higher cell viabilities. Due to the high bioprinting precision across all polymer concentrations, and highest cell viabilities at low polymer concentrations, we utilized the 3 wt% hydrogels for all subsequent studies. To highlight the versatility of our bioprinting approach, we deposited multiple spheroids in close contact into a layered pyramid structure (Fig. 2c i) or as two distinct spheroid populations (pre-labelled with FITC and rhodamine cell masks) into layered rings (Fig. 2cii). These results demonstrate that the shear-thinning and self-healing properties of support hydrogels allow spheroids to be freely moved within the hydrogel for patterning of multi-spheroid structures, without technically challenging intermediate layering and crosslinking steps. There are numerous shear-thinning and self-healing hydrogels that may be useful for this approach; however, these findings indicate that the hydrogel should be evaluated with respect to precision and cell viability prior to use^35^.

**Figure 5:**
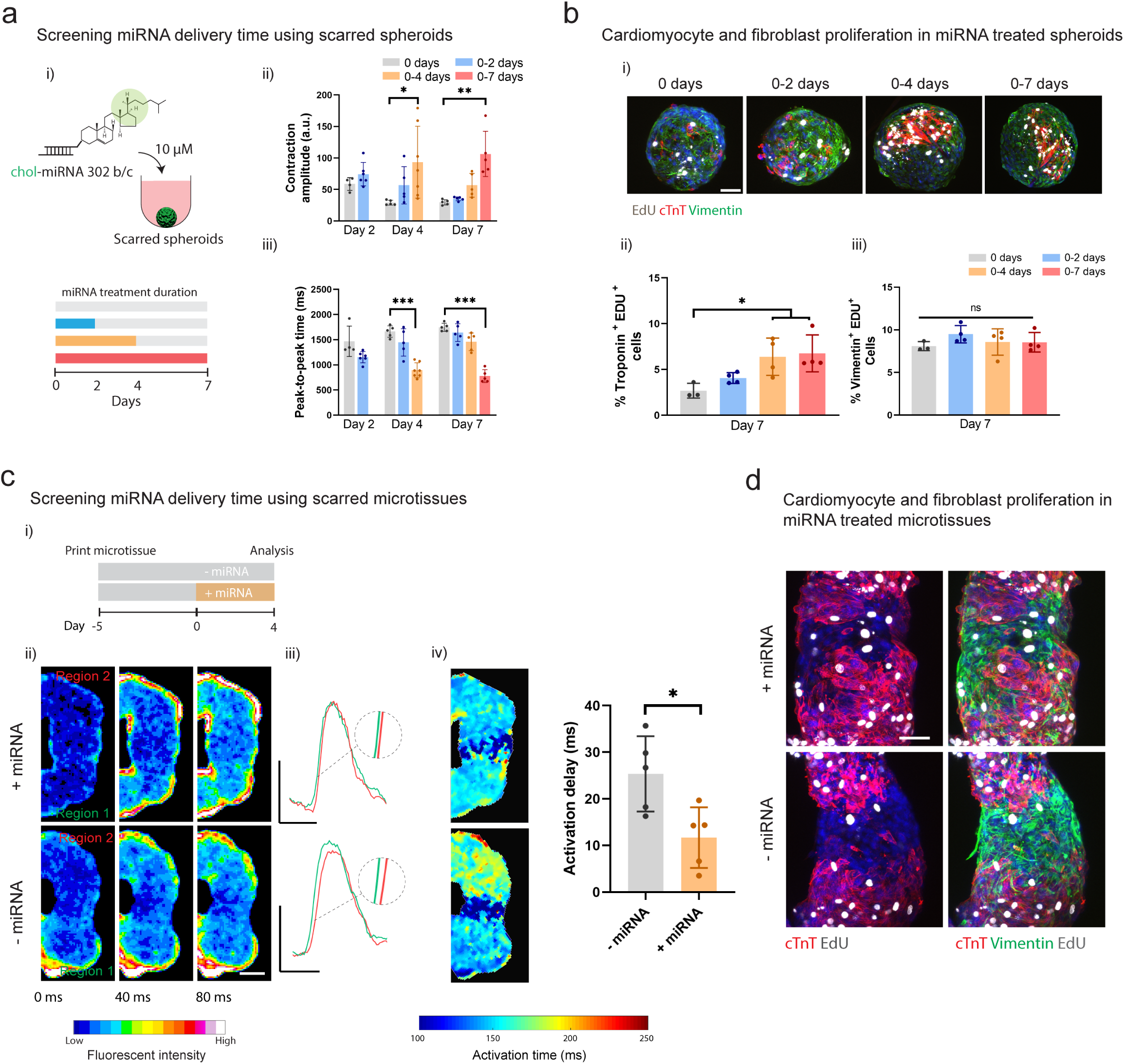
Evaluating miRNA on the behavior of cardiac microtissues. **(a)** (i) Schematic of cholesterol modified miR302 (chol-miRNA 302 b/c) delivery to scarred cardiac spheroids for 0, 0-2, 0-4, and 0-7 days. (ii) Contraction amplitude (a.u) and (iii) peak-to-peak time (ms) within scarred spheroids measured after 2, 4, and 7 days for each treatment period (n=4-7, mean ± s.d, one-way ANOVA *p<0.05 **p<0.01 ***p<0.001 (Day 4; 0 vs. 0-4 days p=0.0138: Day 7; 0 vs. 0-7 days p=0.0045), n=1 donor (B)). **(b)** (i) Immunofluorescence staining for cTnT (red; iPSC-CMs), and vimentin (green; cardiac fibroblasts), and EdU (proliferation marker) in scarred spheroids at day 7 for each treatment condition. (ii) Quantification of cardiomyocyte proliferation (EdU^+^ and cTnT^+^) and (iii) fibroblast proliferation (EdU^+^ and Vimentin^+^) at day 7 for each treatment condition. (n=3-4, mean ± s.d, one-way ANOVA, *p<0.05 (0 vs. 0-4 days p=0.0436, 0 vs. 0-7 days p=0.0258), n.s. (not significant), n=1 donor (B)). Scalebar 50µm **(c)** (i) Experimental outline where scarred microtissues are bioprinted in the support hydrogel as previously described, followed by 4 days treatment with miR302, and subsequent analysis compared to non-treated controls. (ii) Calcium mapping in scarred cardiac microtissues at 9 days (5 days culture in support hydrogel; with and without 4 days miRNA treatment) each image represents a 40 ms interval. Scalebar 100 µm. (iii) Representative calcium traces from regions 1 and 2 in treated and non-treated scarred cardiac microtissues. Scalebars 0.5 ΔF/F_o_ (y), 500 ms (x). (iv) Activation time maps across scarred regions for treated and non-treated scarred cardiac microtissues, and quantification of activation delay (ms) (n=5; two-sided student t-test, *p<0.05 (p=0.0186), n=1 donor (B)). **(d)** Immunofluorescence staining for cTnT (iPSC-CMs), vimentin (CFs), and EdU (proliferation) in scarred cardiac microtissues at day 9 for treated and non-treated scarred cardiac microtissues. Scalebar 50µm.

### 3D bioprinting high-cell density tissue models through directed spheroid fusion

Having established that we could 3D bioprint spheroids at high-resolution inside self-healing support hydrogels, we next explored if directed spheroid fusion could be used to fabricate multi-spheroid microtissues with controlled structures (Fig. 3a i-iii). When spheroids were bioprinted with direct contact, extensive fusion was observed over four days of culture (Fig. 3b i-ii). In addition, when spheroids were bioprinted at a separation distance of 50 µm, comparable levels of fusion were observed (Fig. 3b ii). These distances are well within the measured precision of our method, minimizing any concern that spheroids can be bioprinted with close enough proximity for spheroid fusion. Leveraging these properties, we next deposited 8 spheroids into a ring pattern, which allowed continuous fusion into a microtissue ring after 4 days of culture (Fig. 3c i-ii). This process could be repeated with multiple layers of spheroids to form a microtissue tube (Fig. 3c iii). Importantly, due to the non-covalent and reversible nature of crosslinks in the support hydrogel, it was possible to remove fused microtissues from the support hydrogel for further use (Fig. 3c iii).

When a secondary covalent network was introduced into the support hydrogel to induce more elastic behaviour (Supplementary Fig. 6 i-ii), spheroid fusion was significantly reduced (Supplementary Fig. 6 iii). This highlights the importance of the biophysical properties of the support hydrogel in facilitating spheroid fusion. Spheroids display liquid-like behaviour and undergo coalescence in a similar manner to liquid droplets ^36^, which has been explained using the differential adhesion hypothesis (DAH) where cells rearrange to maximize adhesive cadherin bonding and reduce free energy ^37^. The viscous properties of the shear-thinning hydrogel investigated here allows cells to remodel their surrounding environment in protease-independent mechanisms with local hydrogel yielding at the spheroid-hydrogel interface ^38^, which is reduced in more elastic networks. Such properties should certainly be considered when selecting the appropriate support hydrogel in this approach.

Together these results establish that self-healing hydrogels can be used to support spheroid assembly and fusion into larger 3D microtissues of prescribed structures. The system has a number of advantages when compared to traditional cells-in-gels bioprinting approaches. For example, a significant challenge in the field of biofabrication has been the development of bioinks that meet the rigorous criteria to be processed using extrusion and lithography approaches, while also supporting robust cell differentiation, proliferation and migration ^39^. Here, the biological considerations can be decoupled from the biofabrication process, allowing growth of multi-cellular structures of prescribed 3D patterns. Further, it is possible to fabricate models at organotypic cell densities at early culture times, which is a significant challenge in the bioprinting field, particularly for applications where high levels of cell-cell contact are central to function.

### 3D bioprinting cardiac microtissues for disease modelling applications

There are numerous applications that would benefit from the ability to bioprint high cell density and heterogeneous tissue models. For example, there has been great interest in the engineering of cardiac disease models for personalized medicine applications, including for the screening of drugs for efficacy and toxicity. Indeed, cardiovascular toxicity in therapeutic drug development represents the highest incidence of complications in late-stage clinical development ^40^. Much can be learned from animal models; however, *in vitro* models are more controlled to address fundamental biological questions, can be used with a wide range of imaging and assessment techniques, and can be formulated from patient cells. This has guided the development of engineered human cardiac tissue models that provide controlled environments to study cardiac development and disease along with drug toxicities *in vitro* ^41, 42^. Many technologies are being developed, including self-assembled spheroids/organoids ^6, 43-45^, organ-on-a-chip ^46, 47^, and microtissue platforms^41, 42, 48^. In addition, to better mimic cardiac disease, pathological features are being introduced using hypoxia-induced apoptosis ^6^, mechanical stress ^49^, or varied cardiac cell ratios ^50^. Recent examples have also demonstrated how the introduction of electrical barriers such as holes in healthy cardiac tissue, or fibroblast dense regions can disrupt action potential propagation ^45, 51^.

Despite significant advances in this area, engineering tissue models that accurately replicate the pathophysiology of cardiac disease remains a challenge. Nearly all cardiac diseases involve pathological fibrotic remodeling caused by cardiac fibroblast (CF) proliferation and ECM deposition ^52^. CFs themselves cannot generate action potentials, and excessive ECM production reduces cardiomyocyte-cardiomyocyte (CM) connectivity, leading to reduced tissue compliance and electrical synchronization ^52^. Focal cardiac fibrosis that arises post-MI is a major clinical burden that often leads to a cascade of pathological remodeling and may result in heart failure ^52^. Engineered tissue models that can replicate these pathological features and allow the study of repair mechanisms, could provide a valuable tool for developing therapeutic interventions ^53^.

To address this need, we designed a reductionist model of focal cardiac fibrosis using our spheroid bioprinting technology. To mimic healthy and fibrotic cardiac tissue we fabricated spheroids through the mixing of induced pluripotent stem cell-derived cardiomyocytes (iPSC-CMs) and CFs at varied ratios (4:1 iPSC-CMs to CFs for “healthy”; 1:4 iPSC-CMs to CFs for “scarred”) (Fig. 4a i), which was validated via immunofluorescence staining for cTnT (CM marker) and vimentin (CF marker)(Fig. 4a ii) and marked contractility in healthy spheroids (Fig. 4a iii) at 3 days. Optical mapping of intracellular Ca^2+^ showed robust and synchronized Ca^2+^ flux in healthy spheroids (Supplementary Movie 3), whereas lower levels and only dispersed and sporadic islands of Ca^2+^ flux were observed in scarred spheroids (Supplementary Movie 3) (Fig. 4a iv).

To model focal cardiac fibrosis, we used our bioprinting technology to create heterogenous rings of cardiac tissue through the directed fusion of healthy and scarred spheroids. A ring structure was used to mimic the tissue level electrophysiological properties of a heart chamber, including the possibility of re-entrant arrythmias. To model healthy myocardium, we bioprinted rings of 8 healthy spheroids in direct contact, whereas models of scarred myocardium were bioprinted with rings of 7 healthy spheroids and 1 scarred spheroid (Fig. 4b i). After 5 days of culture, dual staining for cTnT and vimentin confirmed robust fusion between adjacent spheroids, and regions of high fibroblast density were observed in scarred microtissues (Fig. 4b ii). The microtissue rings also contracted in a continuous and synchronized fashion, which could be observed following removal from the support hydrogel (Fig. 4c i, Supplementary Movie 4). Quantification of microtissue contraction revealed reduced contraction amplitudes in scarred microtissues when compared to healthy controls (Fig. 4c ii-iii), mimicking reductions in global contractility that arise following MI.

Next, we used optical Ca^2+^ mapping to assess how fibrotic scars influence electrophysiological properties of microtissues. Healthy microtissues demonstrated synchronized calcium activation propagation across the entire ring, indicating functional electrical coupling between fused spheroids (Fig. 4d i, Supplementary Movie 5). In contrast, wave-like propagation was observed in scarred microtissues (Fig. 4d i, Supplementary Movie 5). Examination of sequential frames from the fluorescent recordings demonstrated limited calcium flux in the scar, and delayed activation between healthy regions on both sides of the scar (Fig. 4d i, Supplementary Movie 5). To visualize activation delays, we plotted calcium activation events in the two healthy regions directly adjacent to the scar, which demonstrated reduced overlap when compared to measurements across the same spheroids in healthy controls (Fig. 4d ii). In addition, activation maps also confirmed increased delays across scarred regions (Fig. 4d iii), which could be quantified as the average activation delay (difference in activation time before and after scar) and was significantly higher than in healthy controls (Fig. 4d ii). Our data shows that bioprinted scars disrupt electrophysiological synchronization in engineered cardiac microtissues, which mimics conduction blocks that can lead to arrythmias following pathological remodeling post-MI ^54^.

Here, we developed a versatile microtissue platform for focal cardiac fibrosis modelling that mimics the two main structural features that arise following scarring (reduced contractile output and electrical synchronization). We chose a relatively simple microtissue design (planar ring, one scar) as *in vitro* models should be designed using a reductionist approach that allows monitoring of variables of interest ^55^. The system is modular and could easily be adopted to model more severe levels of fibrosis by varying CM:CF ratio during spheroid fabrication or through bioprinting multiple and/or larger scars. The ability to create healthy and diseased tissue in a single microtissue also offers the opportunity to probe borderzone interactions and study aspects of cardiac repair *in vitro*.

In contrast to our approach, traditional extrusion-based bioprinting approaches can be challenging to implement for cardiac models. Functional cardiac muscle requires high levels of cell-cell contact to support rapid action potential propagation and synchronized contraction. Fibrillar hydrogels such as collagen and fibrin are compatible with cardiac tissue engineering as the cells can rapidly contract the matrix to form high cell density tissues ^48^; however, these softer hydrogels are challenging to process using biofabrication technologies ^39^. The organotypic cell densities that can be achieved using our self-assembled spheroid approach also supports high levels of CM-CM contact, which is analogous to healthy cardiac tissue. It should be noted that spheroid-based microtissues developed in our system lack the anisotropic tissue alignment present in mature myocardium and do not include vascular networks that are highly abundant in native myocardium; however, future efforts could advance the platform with additional control over spheroid anisotropy and the incorporation of endothelial cells, as has been previously described for cardiac spheroids ^44^.

### Evaluating the behavior of cardiac microtissues in response to miRNA therapeutics

Having established a 3D bioprinted cardiac fibrosis model, we next looked to utilize the system to study microRNA (miRNA) therapeutics for cardiac repair. Following cell death within the ischemic environment after MI, cardiac tissue has limited capacity for self-repair in the adult, particularly due to the non-proliferative nature of CMs. Recent pre-clinical evidence indicates that Hippo pathway inhibiting miRNAs can stimulate CM proliferation ^34, 56, 57^. For example, the delivery of Hippo inhibiting miR302 b/c mimics stimulate cardiomyocyte proliferation and cardiac repair following MI in mice^34^. However, in a more recent porcine MI model, prolonged expression of Hippo inhibiting miRNA-199a through viral transfection resulted in sudden cardiac failure in 70% of the treated pigs ^58^. This was attributed to excessive cardiomyocyte dedifferentiation and proliferation that produced islands of non-contractile tissue and fatal arrythmias. These results indicate that cardiac regeneration can be stimulated through miRNA therapies targeting cardiomyocyte proliferation; however, their timing needs to be precisely controlled to avoid detrimental outcomes. Our bioprinted cardiac microtissues present an opportunity to evaluate how the extent of miR302 b/c induced Hippo inhibition (treatment duration) influences cardiomyocyte proliferation and cardiac repair (e.g., contractility, calcium propagation).

First, we screened a range of miRNA treatment durations (0, 0-2, 0-4, and 0-7 days) using our scarred cardiac spheroids (Fig. 5a i). To assess the transient effects of miRNA treatment, we performed sequential contraction imaging at 2, 4, and 7 days. Significant increases in contraction amplitude were observed with constant treatment for 4 and 7 days when compared to controls (Fig. 5a ii). Notably, contraction values increased approximately 3-fold for these conditions and approached ∼20% of the contraction amplitudes present in healthy spheroids. Interestingly, the speed of contraction also increased significantly when compared to controls (approximately 2X faster) when miRNA was still present in the media (0-4 days treatment at 4 days, 0-7 days treatment at 7 days) (Fig. 5aiii). We also assessed how miRNA treatment influenced the contractile properties of healthy spheroids (Supplementary Fig. 7a). No significant changes in contraction amplitude were observed in health spheroids (Supplementary Fig. 7a ii), with an increase in contraction speed at earlier timepoints only (0-2 days treatment at 2 days) (Supplementary Fig. 7a iii).

To determine if miRNA treatment promoted CM and CF proliferation in scarred spheroids, we performed EdU labelling (a proliferation marker) along with co-staining for cTnT and vimentin staining at 7 days (Fig. 5b i). Interestingly, islands of cTnT^+^EdU^+^ were present, suggesting proliferation and clonal expansion (Fig. 5b i), particularly with longer treatment durations (0-4 days, 0-7 days) (Fig. 5b ii). No increases in the number of Vimentin^+^ Edu^+^ cells were observed, indicating that miRNA treatment was primarily influencing CM proliferation (Fig. 5b iii). Similar trends were also observed in healthy spheroids, with increases in the number of cTnT^+^ and Edu^+^ cells observed for all treatment conditions when compared to controls (Supplementary Fig. 7b i-ii) and limited effects on CF proliferation (Supplementary Fig. 7b iii).

Having established that miR302b/c mimics stimulate CM proliferation and increase the functional properties of scarred spheroids, we next assessed if these improvements could translate into tissue level repair using our scarred microtissue model. To do this, scarred microtissues were bioprinted as previously described, followed by treatment with miRNA for 4 days (Fig. 5c i). We chose 4 days treatment, as further improvements in contraction or CM proliferation were not observed with 7 days of continuous treatment. At 4 days treatment, we assessed tissue-level electrophysiological properties through calcium mapping (Fig. 5c ii). Further corroborating the improvements found with single spheroids, miRNA treatment significantly reduced activation delays present in scarred microtissues when compared to untreated controls (Fig. 5c iii). Finally, histological staining at the scar interface allowed us to visualize tissue remodeling in response to miRNA treatment (Fig. 5d).

Engineered cardiac tissues are being widely used as models to study drug toxicities and efficacy *in vitro* ^6, 42-44, 59^. Here, we developed a fibrosis model that enabled the study of therapeutics for cardiac repair, including with functional outcomes to assess changes in contraction and electrophysiological properties. The introduction of scars adjacent to healthy tissue allowed us to study tissue-level electrophysiological properties and our results demonstrate that these fibrotic regions are responsive to therapeutics. In our model, miR302 b/c mimics enhanced iPSC-CM proliferation, resulting in enhanced contractility in scarred spheroids and improved electrophysiological integration. Interestingly, miRNA treatment did not enhance cardiac fibroblast proliferation which is an important consideration when designing miRNA therapeutics for cardiac repair and corroborates previous 2D cultures of iPSC-CMs and CFs with miRNA treatment ^60^. Our findings highlight how 3D bioprinting technology can be used to design personalized *in vitro* disease models that provide comparable functional outputs to pre-clinical animal models, while keeping investigations simple, minimizing costs, and in formats that support a wide variety of imaging and assessment techniques.

## Conclusion

Advances in bioprinting technologies have enabled the development of *in vitro* models and implantable constructs that better mimic the complexity of native tissues and organs. Many of these technologies rely on processing cell suspensions embedded in hydrogels, making it challenging to engineer tissues with native tissue-like cell-densities and heterogeneity, and resulting functions. Here, we developed a new process that enables the 3D bioprinting of high-cell density tissue models, with precise control over microtissue structure and local heterogeneity. This is enabled by the application of self-healing hydrogels that support the bioassembly and fusion of bioprinted spheroids in 3D space to form continuous cell-dense tissue models. We demonstrate the potential of the approach by bioprinting cardiac microtissue disease models that recapitulate pathological scarring features that arise post-MI, and using readouts for cardiac function (contraction, electrophysiological synchronization) we were able to probe miRNA therapeutics for repair. The approach is highly generalizable and could be implemented with a wide range of spheroid and organoid systems, which opens-up many new opportunities in 3D bioprinting of precision models to mimic diseases and for drug screening.

## Methods

### Hydrogel preparation

HA macromers (adamantane (Ad)-HA, cyclodextrin (Cd)-HA) were synthesized as previously described ^32, 61^. Briefly, the tetrabutylammonium salt of sodium hyaluronate (Lifecore, 64 kDa) (HA-TBA) was prepared using Dowex 50W proton exchange resin. Ad was coupled to the tetrabutylammonium salt of HA (HA-TBA) to obtain Ad-HA through 4-dimethylaminopyridine (DMAP)/ditert-butyl dicarbonate (BOC_2_O)-mediated esterification. For CD-HA, aminated β-CD was coupled to HA-TBA via amidation in the presence of benzotriazole-1-yl-oxy-tris-(dimethylamino)-phosphonium hexafluorophosphate (BOP). Products were dialyzed, frozen, and lyophilized. Next, ^1^HNMR (DMX 360 MHz, Bruker) was used to determine that Ad-HA had ∼19.5% of HA repeat units modified with Ad (Supplementary Fig. 2a) and CD-HA contained ∼18.5% of HA repeat units modified with β-CD (Supplementary Fig. 2b) AdNorHA was synthesized as previously described ^62^. To form support hydrogels, Ad-HA and CD-HA macromers were separately dissolved in PBS and then mixed at a 1:1 Ad:CD ratio and final polymer concentrations of 3, 5, and 7 wt%. To introduce secondary covalent crosslinking into the hydrogel, Ad-HA was replaced with AdNorHA and supplemented with 1mM DL-dithiothreitol (DTT) and 0.05 wt% photoinitiator (lithium phenyl-2,4,6-trimethylbenzoylphosphinate (LAP), Colorado Photopolymer Solutions (Boulder, CO)). Crosslinking was performed with the application of visible light (2 mW cm^-2^ intensity, 400-500 nm wavelength, 3 min exposure).

### Cell culture and spheroid formation

Human MSCs (hMSCs) were isolated from fresh unprocessed bone marrow from human donors (Lonza) as previously described ^63^. Briefly, diluted bone marrow (1:4 with PBS) was separated with Ficoll density gradient centrifugation (800 RCF, 20 min). Mononuclear cells were collected from the liquid interface, plated on tissue culture plastic, cultured in alpha-modified essential medium (α-MEM, 10% FBS, 1% penicillin/streptomycin, 5 ng ml^−1^ basic fibroblast growth factor) at 37 °C/5% CO_2_ until 80% confluency of the colonies, followed by storage in liquid nitrogen (95% FBS, 5% DMSO). Next, hMSCs were expanded in standard growth media (α-MEM, 10% FBS, 1% penicillin/ streptomycin) for one passage before spheroid formation. hMSC spheroids were prepared by centrifugation in ultra-low attachment 96-well round-bottom plates (Corning 7007, 500G). Two spheroid sizes were prepared by adding either 5,000 or 10,000 hMSCs/well, which resulted in average spheroid diameters of ∼200µm and ∼400µm (Supplementary Fig. 1c). For visualization, hMSC spheroids were labelled with Cell Tracker™ fluorescent probes (Green-CMFDA, Red CMPTX, 5µM).

Human iPSC-CMs were purchased from Fujifilm cellular dynamics (icell 01434 (Donor A) & 11713 (Donor B), purity >97%). Human cardiac fibroblasts (CFs) were purchased from Promocell and expanded in α-MEM (10% FBS, 1% penicillin/streptomycin, 5 ng ml^−1^ basic fibroblast growth factor) at 37 °C/5% CO_2_ until 80% confluency. Cardiac spheroids were formed by mixing iPSC-CMs and CFs at different ratios (4:1 for “healthy” and 1:4 for “scarred”, 5000 cells total) in ultra-low attachment 96-well round-bottom plates. For the first 48 hours cardiac spheroids were cultured in iCell plating medium (Cellular dynamics) to optimize post-thawing survival of the iPSC-CMs. After 48 hours, cardiac spheroids were maintained in iCell maintenance media (Cellular dynamics). Approximately 24 hours post-thawing the iPSC-CM and HCF co-cultures had aggregated into a single spheroid with synchronized contractions observed after 24-48 hours. For all further experiments, cardiac spheroids were used at 96 hours post-thawing.

### Rheological characterization

The viscoelastic properties of the support hydrogels were characterized using shear rheometry (cone and plate geometry with (0.995°) cone angle, 27 µm gap, TA Instruments, AR2000) at 25 °C. To assess shear-thinning properties, viscosity was measured as a function of shear rate (0 to 50 s^-1^). Shear-recovery was assessed by applying low (0.5%) and high strains (100%) periodically at 10 Hz. Strain yielding behaviour was assessed using strain sweeps (0-500%, 10 Hz).

### 3D bioprinting setup

Custom molds for 3D bioprinting were drawn using computer-aided design (CAD) software (Solidworks) and then 3D bioprinted from Accura SL 5530 (Proto Labs). PDMS molds were then cast against the molds and bonded to a glass coverslip (Supplementary Fig. 1a). The mold system consisted of two reservoirs (200µl volume) connected by a narrow channel (750µm width) (Supplementary Fig. 1a). The first reservoir was filled with cell culture media and spheroids and the second reservoir was filled with the support hydrogel (Supplementary Fig. 1a). The connecting channel was designed to prevent flow between the two reservoirs. Next, the PDMS mold containing spheroids and support hydrogel was placed in a live cell incubation chamber (37°, 5% CO_2_) and spinning disk confocal (Nikon TE-2000) was used to visualize spheroids during bioprinting (Supplementary Fig. 1b). To bioprint spheroids in the support hydrogel, a micromanipulator with XYZ control (Transferman®, Eppendorf) containing custom pulled micropipette tips (Ø 100µm) was used to vacuum aspirate the spheroids, followed by transfer via the connecting channel into the support hydrogel (500 µm/s), and then deposition with vacuum removal (Supplementary Figs. 1a,b, Supplementary Movie 1).

To visualize the support hydrogel during bioprinting, green-fluorescent polystyrene beads were added at a 1:200 dilution (nominal diameter ∼1 µm, emission/excitation 441/485 nm, Fluoresbrite®). Spheroids were translated through the hydrogel at a constant speed (400 µm/s) and microparticle motion was tracked using fluorescent videos (20 HZ) (Supplementary Movie 2). To measure bead displacements, image frames were isolated from the videos every 50 frames, and the relative bead motion between frames was used to produce vector and displacements maps using the particle image velocimetry (PIV) plugin (Image J) (Supplementary Fig. 3).

### 3D bioprinting accuracy measurements

To quantify bioprinting precision in the XY plane, a partially transparent target was superimposed on the bottom glass surface of the bioprinting mold (Supplementary Fig. 4a) and a spheroid was bioprinted directly above the target. Next, the bioprinting nozzle was removed, and the post-printing drift was measured after 5 minutes by measuring the distance between the center of the spheroid and the target. To quantify bioprinting precision in the Z plane, a spheroid was deposited in the support hydrogel and the z-position was monitored over 24 hours using brightfield videos (Supplementary Fig. 4b).

### Live/dead staining

For viability analysis, spheroids encapsulated in hydrogels were stained using a Live/Dead cell-viability assay (Invitrogen) according to the manufacturer’s instructions. Viability was quantified from confocal stacks (250 µm thick) acquired using a Leica SP5 II confocal microscope. The live cell area was reported as the ratio of calcein-AM stained area to the total spheroid area (sum of calcein-AM stained area and ethidium homodimer-1 stained area).

### Microtissue 3D bioprinting and spheroid fusion measurements

To 3D bioprint microtissues, 8 MSC spheroids were bioprinted into the support hydrogel in a circular ring with direct contact between adjacent spheroids. Molds containing bioprinted spheroids in the support hydrogel were cultured at 37 °C/5% CO_2_ and the media reservoir was refreshed every two days. For all hMSC spheroid experiments, α-MEM (10% FBS, 1% penicillin/streptomycin, 5 ng ml^−1^ basic fibroblast growth factor) was used. To track spheroid fusion dynamics hMSC spheroids were labelled with Cell Tracker™ fluorescent probes (Green-CMFDA) and fluorescent images were taken using a spinning disk confocal (Nikon TE-2000). The spheroid fusion index was calculated as described (Supplementary Fig. 6). Microtissues were then removed from the support hydrogel through gentle aspiration using a microcapillary tip, followed by transfer back into the media reservoir.

### 3D bioprinting cardiac microtissues and contraction imaging

To 3D bioprint healthy cardiac microtissues, 8 healthy spheroids were bioprinted into the support hydrogel in a circular ring as described above. Scarred microtissues were bioprinted by including 1 scarred spheroid and 7 healthy spheroids into a circular ring. For all cardiac spheroid experiments, samples were maintained in iCell maintenance media (Cellular dynamics) at 37°/5% CO_2_. After 5 days of culture and spheroid fusion, the bioprinted cardiac rings were removed from the support hydrogels using aspiration and transferred back into the media reservoir. A thin layer of agarose (3 wt%) was added to the bottom surface to prevent attachment of the microtissues to the coverslip. Microtissue contraction was recorded at 37°/5% CO_2_ in iCell maintenance media using a Nikon TE2000 (4x lens, 20 HZ). The software package musclemotion was used for all contraction analysis ^64^.

### Calcium imaging and analysis

To assess calcium activation, microtissues were treated with intracellular calcium indicator (Cal 520, AAT Bioquest, 4µM) for 90 minutes. Pluronic (0.04%) was included to enhance cellular uptake and all staining solutions were prepared in iCell maintenance media. Following calcium loading, the staining solution was replaced with maintenance media and fluorescence images of calcium concentration were acquired with a Nikon Eclipse TE2000U microscope fitted with a spinning disk confocal (CSU-10b, Solamere Technologies), a CCD camera (Photometric Cool-Snap HQ^2^, BioVision, Exton PA) (4×4 binning to 174 × 130 pixels), 488-nm excitation laser (Prairie SFC), and a Nikon 4X Plan Apo objective (N.A. = 0.2). Exposure time was set to 15 ms, and images were streamed at 50 Hz frame rate for at least 1000 frames.

The high-throughput and open-source software package ElectoMap was used to perform quantitative measurements on the acquired fluorescent calcium videos ^65^. Activation maps were produced using the isochronal module using a spatial (2 pixels, sigma 1.5) and temporal (Savitsky-Goaly) filter, and the activation measure was defined as the activation midpoint. Activation delays were then calculated by measuring differences in local activation time across the microtissues. For example, in scarred microtissues the activation delay was calculated by measuring the difference in activation time in the two healthy regions directly adjacent to the scarred region. In healthy microtissues, the activation time was averaged in six evenly space regions across the microtissues, and the activation delay was calculated by measuring the average activation delay across each of these regions. All calcium traces presented represent the ensemble averaged signal and were produced using Electromap.

### Immunofluorescent staining and analysis

Microtissues were washed 3X times with PBS and then fixed overnight in 10% formalin at 4 °C. Following 3X washed with PBS, fixed microtissues were incubated for 4 h at 4 °C in permeabilization solution containing 0.2% Triton X-100 in PBS supplemented with 2% BSA, 320 mM sucrose and 6 mM magnesium chloride. Before immunostaining, microtissues were blocked in PBS containing 2% BSA for 2 hours. Next, microtissues were stained with primary antibodies for cardiac troponin-t (cTnT) (Thermofischer MA5-12960, 1:200) and vimentin (MA5-11883; 1:200) to visualize cardiomyocytes and fibroblasts, respectively. Primary antibodies were diluted in PBS containing 2% BSA and hydrogels were stained at 4 °C for 72 hours. After three PBS washes, the secondary antibodies Alexa Fluor-488/594/647 IgG H&L (1:200; Abcam ab150113, ab150080, ab150075) were added at 4 °C for 72 hours. Finally, samples were washed 3X in PBS followed by DAPI staining (1:2000; Invitrogen D1306, in PBS) for 30 min at room temperature. A Leica SP5 confocal was used to acquire z-stack images.

### miRNA treatments and proliferation analysis

Cholesterol modified miR-302b/c was purchased from GE Dharmacon. Their sequences are as follows.

<u>Mus musculus miR-302b (miR-302b):</u>

-5′ -

ACUUUAACAUGGGAAUGCUUUCU-

3′ (guide)

-3′ -

GAUGAUUUUGUACCUUCGUGAAU-

chol-5′ (passenger)

<u>Mus musculus miR-302c (miR-302c):</u>

5′ -GCUUUAACAUGGGGUUACCUGC-

3′ (guide)

3′ -GGUGACUUUGUACCUUCGUGAA-

chol-5′ (passenger)

For all treatments, miR-302b/c mimics were diluted in iCell maintenance media at 10µM (each) and media was refreshed every 48 hours. To assess cardiomyocyte and fibroblast proliferation during miRNA treatment, a Click-iT® EdU Protocol (C10337 Thermofisher) was used. In this assay, the modified thymidine analog EdU (5-ethynyl-2’-deoxyuridine, a nucleoside analog of thymidine) is incorporated into newly synthesized DNA and fluorescently labeled with Alexa Fluor 488 dye. During miRNA treatment of cardiac microtissues, 5 µM EdU labeling solution was added to the culture media along with the miR-302b/c mimics. Then following fixing and immunofluorescence staining described above, EdU incorporated nuclei were visualized through a click reaction with Alexa Fluor 488 azide. Note for EdU staining with Alexa Fluor 488, the secondary antibodies Alexa Fluor-594/647 IgG H&L (1:200; Abcam ab150080, ab150075) were used to label the cTnT and vimentin primary antibodies. A Leica SP5 confocal was used to acquire z-stack images (300 µm thick, 3µm slices) and the number of EdU^+^/cTnT^+^ and EdU^+^/Vimentin^+^ cells were manually counted using image-J to quantify iPSC-CM and CF proliferation. The percentage of proliferating cells were then normalized to total cell number by quantifying the total number of DAPI stained nuclei.

### Statistical analysis

GraphPad Prism 8 software was used for all statistical analyses. Statistical comparisons between two experimental groups were performed using two-tailed Student’s t-tests and comparisons among more groups were performed using a one-way ANOVA with Bonferroni post hoc testing. All measurements were taken from distinct samples with no repeated measures. Sample distribution was assumed normal with equal variance.

## Acknowledgements

The authors would like to acknowledge assistance from Dayo Adewole and Professor D. Kacy Cullen for assistance with calcium imaging and Gladys Gray for assistance with confocal imaging. This work was made possible by financial support from the American Heart Association through a postdoctoral fellowship (A.C.D.) (20POST35210923), the National Institutes of Health (F32 DK117568), and the National Science Foundation supported Center for Engineering MechanoBiology STC (CMMI: 15-48571).

## Data availability

All data supporting the results in this study are available within the article and its supplementary information. The range of raw datasets acquired and analyzed are available from the corresponding author on reasonable request.

## Author contributions

A.C.D. and J.A.B. conceived the ideas and designed the experiments. A.C.D. and M.D.D. conducted the experiments and analyzed the data. A.C.D., M.D.D., and J.A.B. interpreted the data and wrote the manuscript. All authors have given approval to the final version of the manuscript.

