## Supplementary figures for "3D bioprinting of high cell-density heterogeneous tissue models through spheroid fusion within self-healing hydrogels"

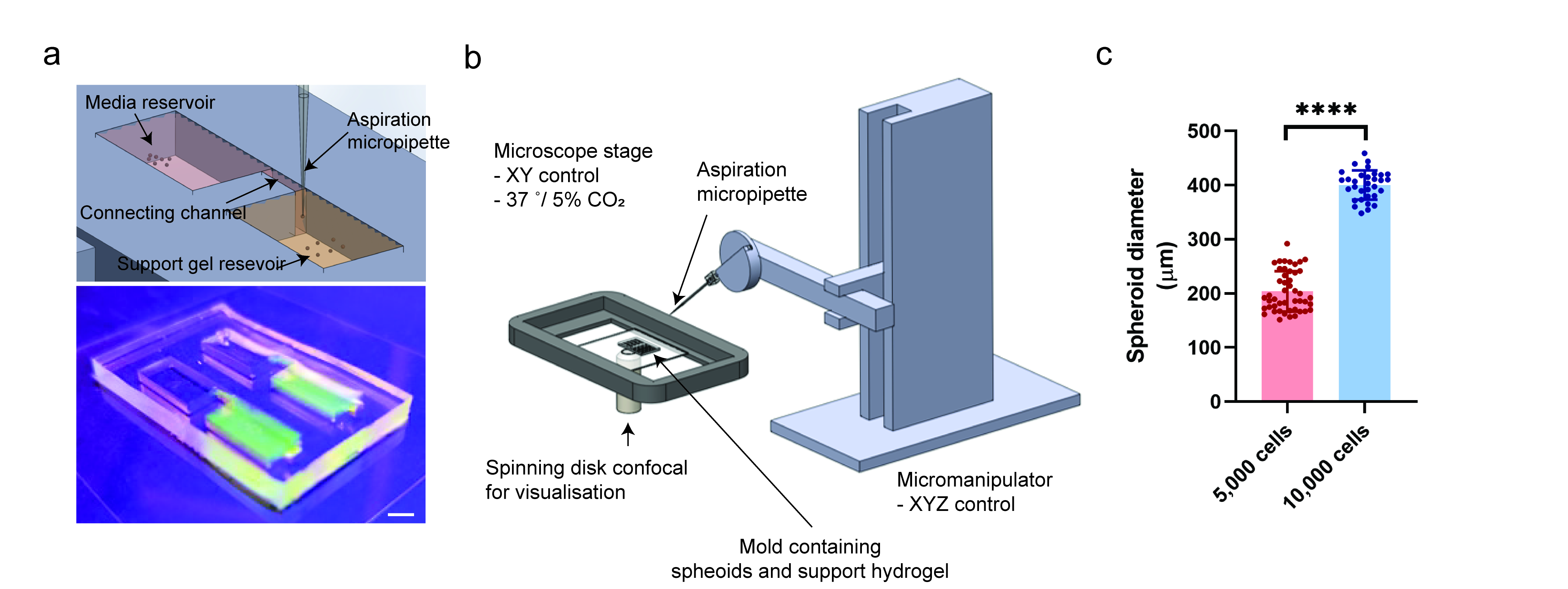

**Supplementary Figure 1: 3D bioprinting approach and spheroid formation. (a)** Schematic of the 3D bioprinting molds used to create a media reservoir and a support hydrogel reservoir connected with a narrow channel, scale bar 7 mm.  **(b)** Schematic of the 3D bioprinting setup composed of a robotic micromanipulator (XYZ spatial control) with a microcapillary needle for spheroid aspiration, a microscope stage (XY spatial control) with an environmental chamber (37°C, 5% CO_2_), and a spinning disk confocal for live brightfield and fluorescent imaging during bioprinting. **(c)** Average spheroid diameter (µm) 24 hours after seeding of either 5,000 or 10,000 human MSCs in ultra-low attachment 96 well round bottom plates. (n>32, mean ± s.d, two-sided student t-test **** p<0.0001).

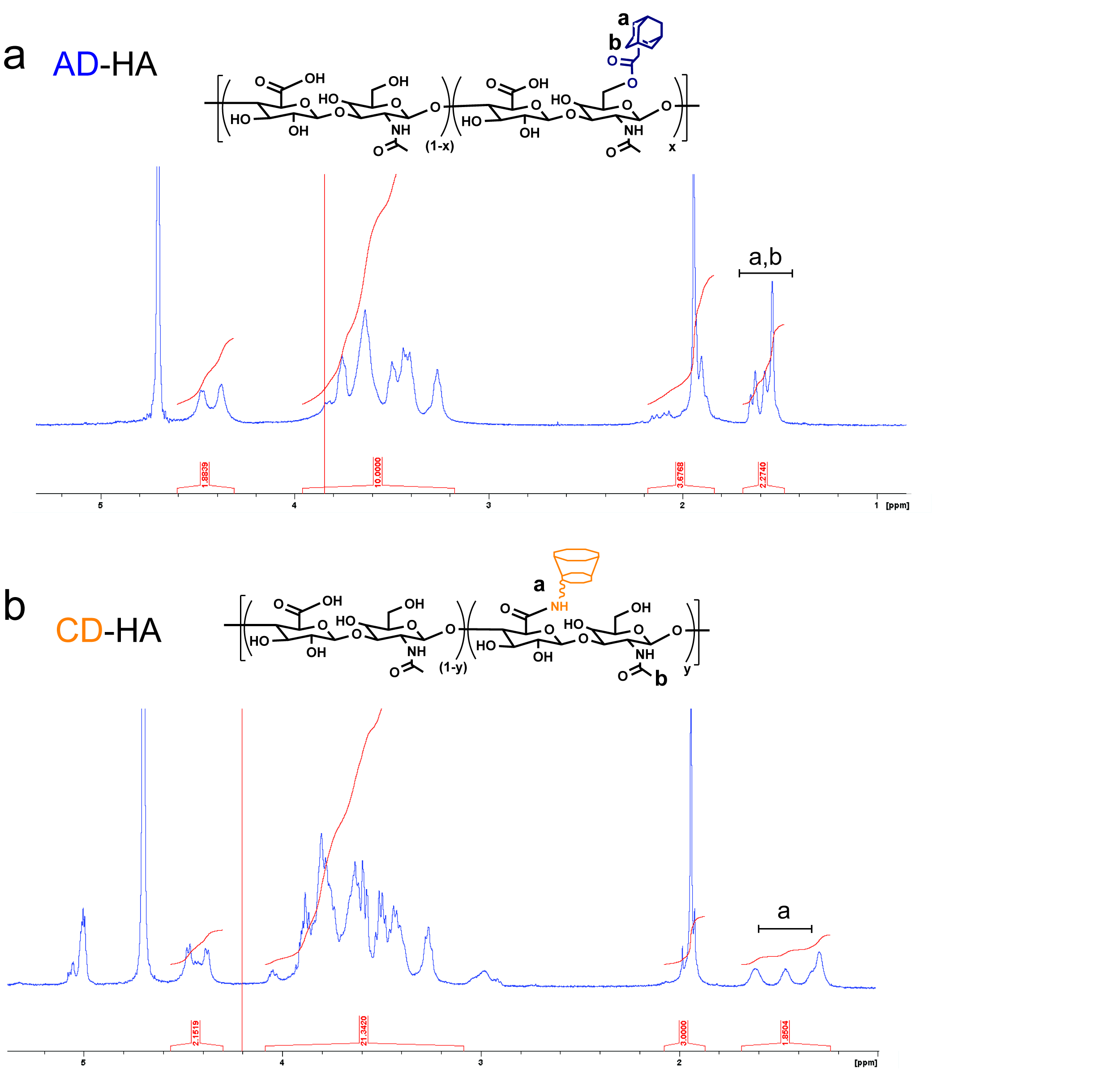

**Supplementary Figure 2: Characterization of synthesized HA polymers.** **(a)** ^1^H NMR spectrum of adamantane modified hyaluronic acid (Ad-HA); the degree of adamantane functionalization (~18.9 %) relative to HA disaccharides (δ = 3.26 – 3.95, 10 H) is determined by integration of adamantane ethyl multiplet (δ = 1.56 - 1.87, 12 H, 2.274). **(b)** ^1^H NMR spectrum of β-cyclodextrin-modified hyaluronic acid (CD-HA); the degree of β-cyclodextrin functionalization (~15.4 %) relative to the HA methyl singlet (δ = 2.1, 3 H) is determined by integration of the hexane linker (δ = 1.16 - 1.65, 12 H, 1.8504).

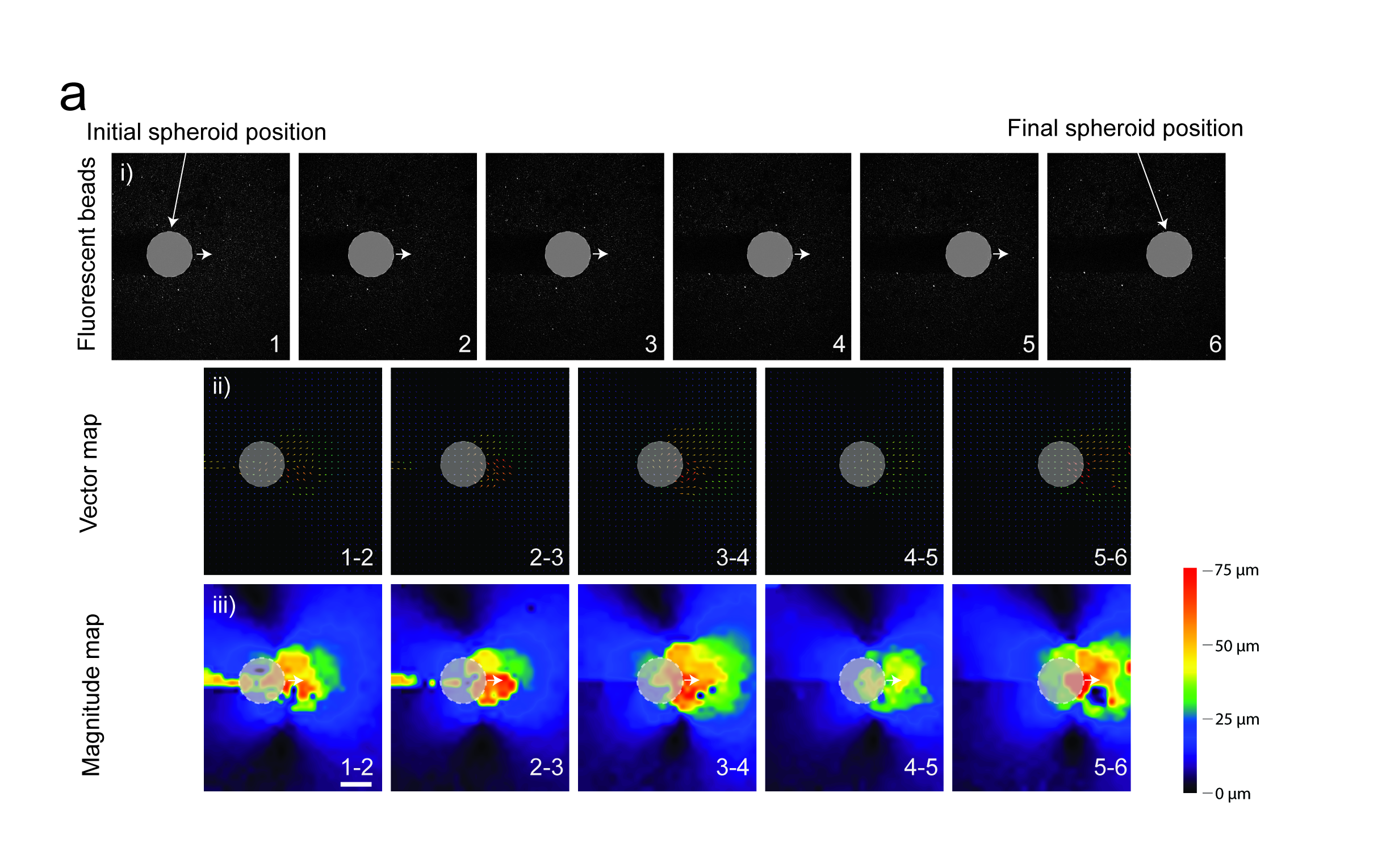

**Supplementary Figure 3: Tracking support hydrogel motion during spheroid bioprinting.** (i) Sequential images showing the motion of fluorescent beads within the support hydrogel during spheroid translation from left to right (1µm Ø fluorescent beads (white), 1:200 dilution). (ii) Particle image velocimetry (PIV) vector plots and (iii) particle image velocimetry (PIV) magnitude plots demonstrating relative bead motion between frames presented in the top panel, e.g. 1-2 represents the bead motion between images 1 and 2. Scalebar 200 µm.

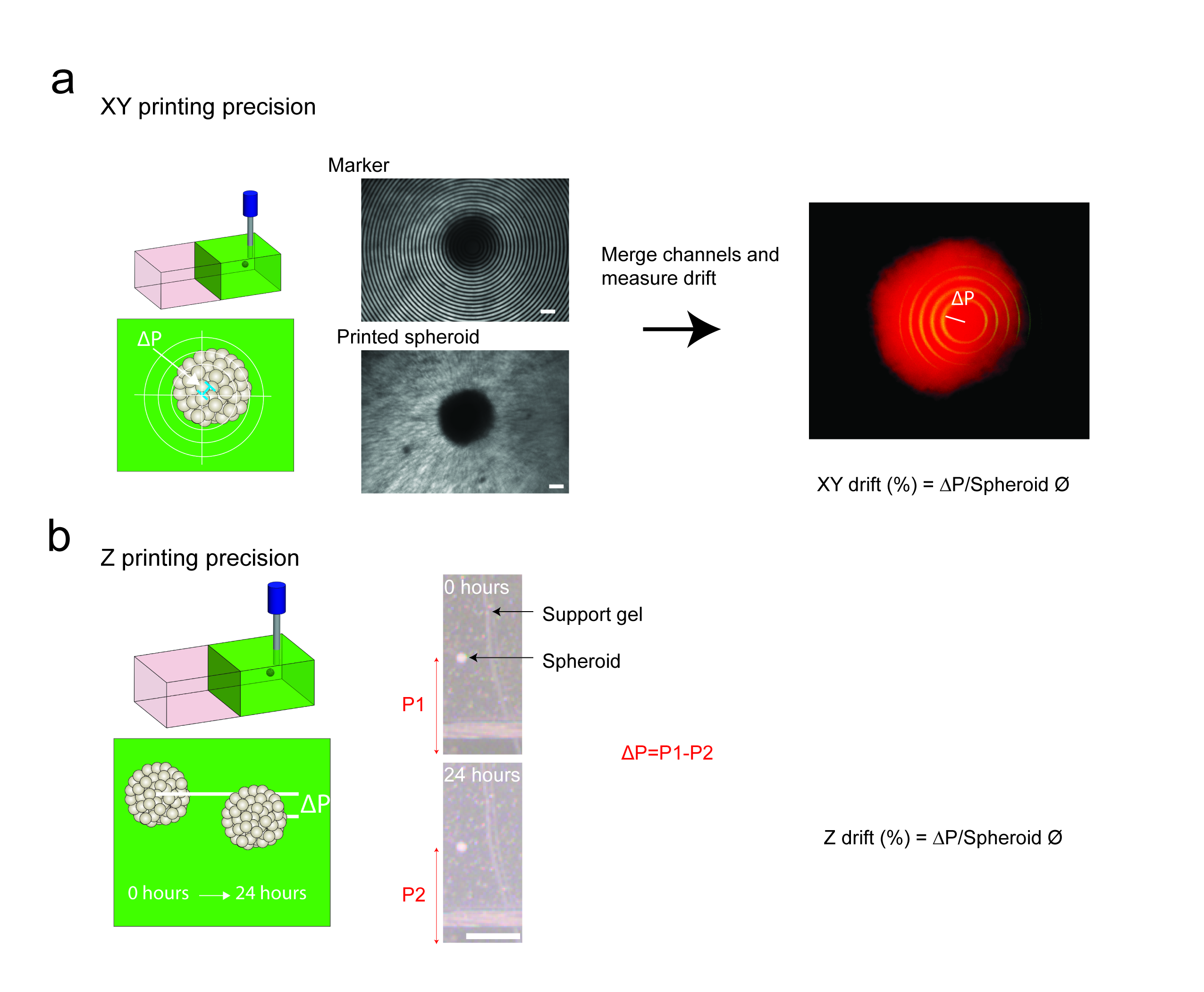

**Supplementary Figure 4: 3D bioprinting precision measurements. (a)** XY bioprinting precision method where a spheroid is deposited directly above a crosshatch marker and the post-printing spheroid drift (5 minutes after removing nozzle) is measured from the center of the spheroid. Scalebar 100µm. XY drift % = ΔP_xy_/spheroid diameter. **(b)** Z bioprinting precision method where a bioprinted spheroid’s z position is tracked in the support hydrogel over 24 hours. Scalebar 1mm. Z drift % = ΔP_z_/spheroid diameter.

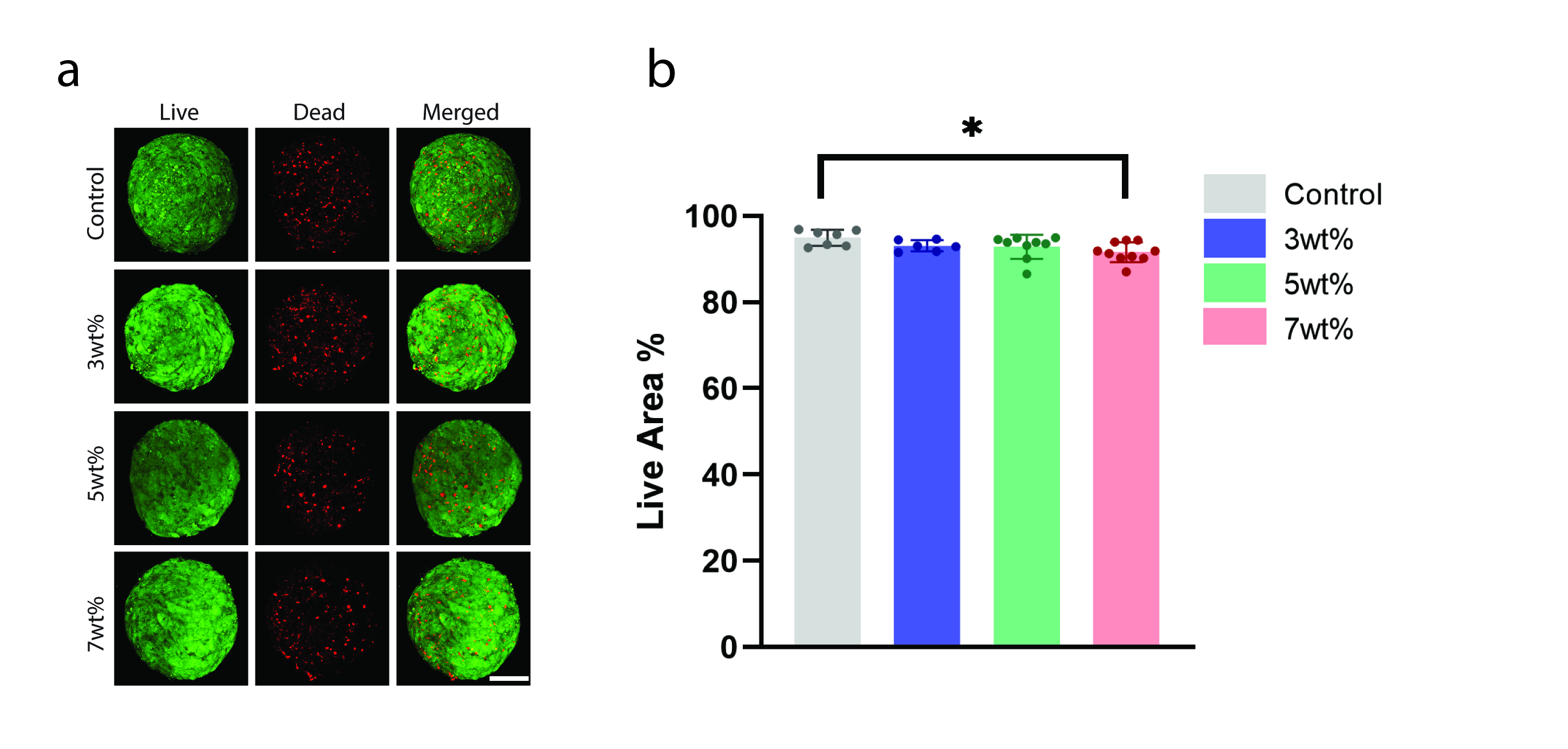

**Supplementary Figure 5: Post-printing cell viability. (a)** Spheroid live/dead staining in 3, 5, and 7 wt% support hydrogels compared to non-printed controls 24 hours post-printing (note: control spheroids were added to the media reservoir, but not bioprinted into a support hydrogel). Scalebar 100 µm **(b)** Quantification of spheroid live area for each condition. (n=6-10, mean ± s.d, one-way ANOVA *p<0.05 (Control vs. 7wt% p=0.0248).

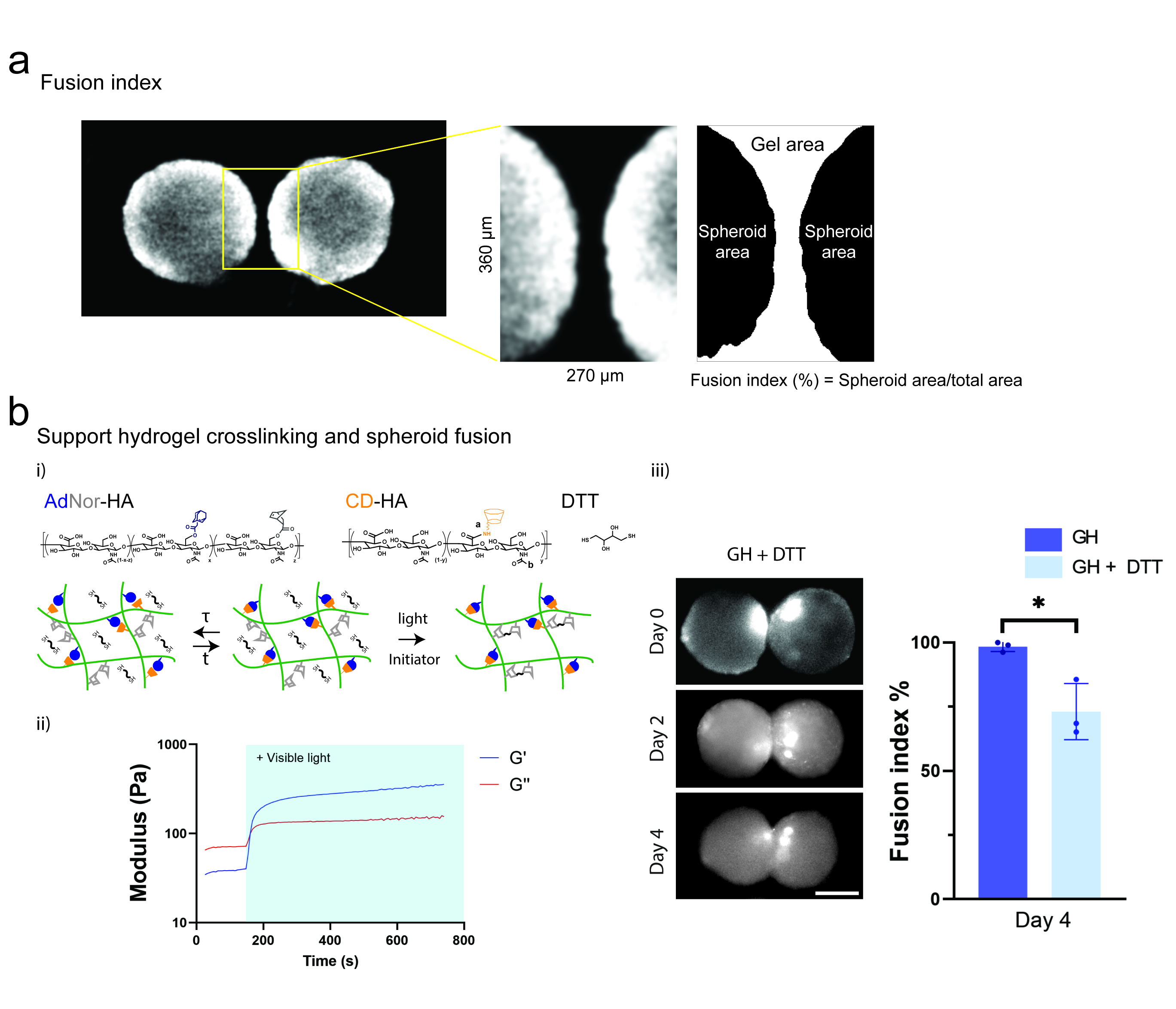

**Supplementary Figure 6: Spheroid fusion dynamics. (a)** Spheroid fusion measurements where the spheroid area (fluorescently labelled) and gel area (non-fluorescently labelled) between two adjacent spheroids (360 µm X 270 µm box) is quantified and used to determine the fusion index. Fusion index % = Spheroid area/ total area. Scalebar 200 µm. **(b)** (i) Secondary crosslinking of the guest-host support hydrogel through a thiol-ene reaction between norbornene groups and d-thiol crosslinkers (DTT, non-degradable) in the presence of visible light. (ii) Rheological characterization of support hydrogel in response to secondary stabilization, time sweep (1 Hz, 1% strain). This indicates that the support hydrogel initially displays viscous behaviour, with a transition towards more elastic properties (increase in G’) after thiol-ene crosslinking of norbornene groups with DTT (1mM DTT, 0.05% LAP photoinitiator, 2 mW cm^-2^ intensity 400-500 nm wavelength)**.** (iii) Spheroid fusion over 4 days with and without secondary crosslinking in the presence of DTT. (n=3, mean ± s.d, two-sided student t-test *p<0.05 (p=0.017)). Scalebar 200 µm.

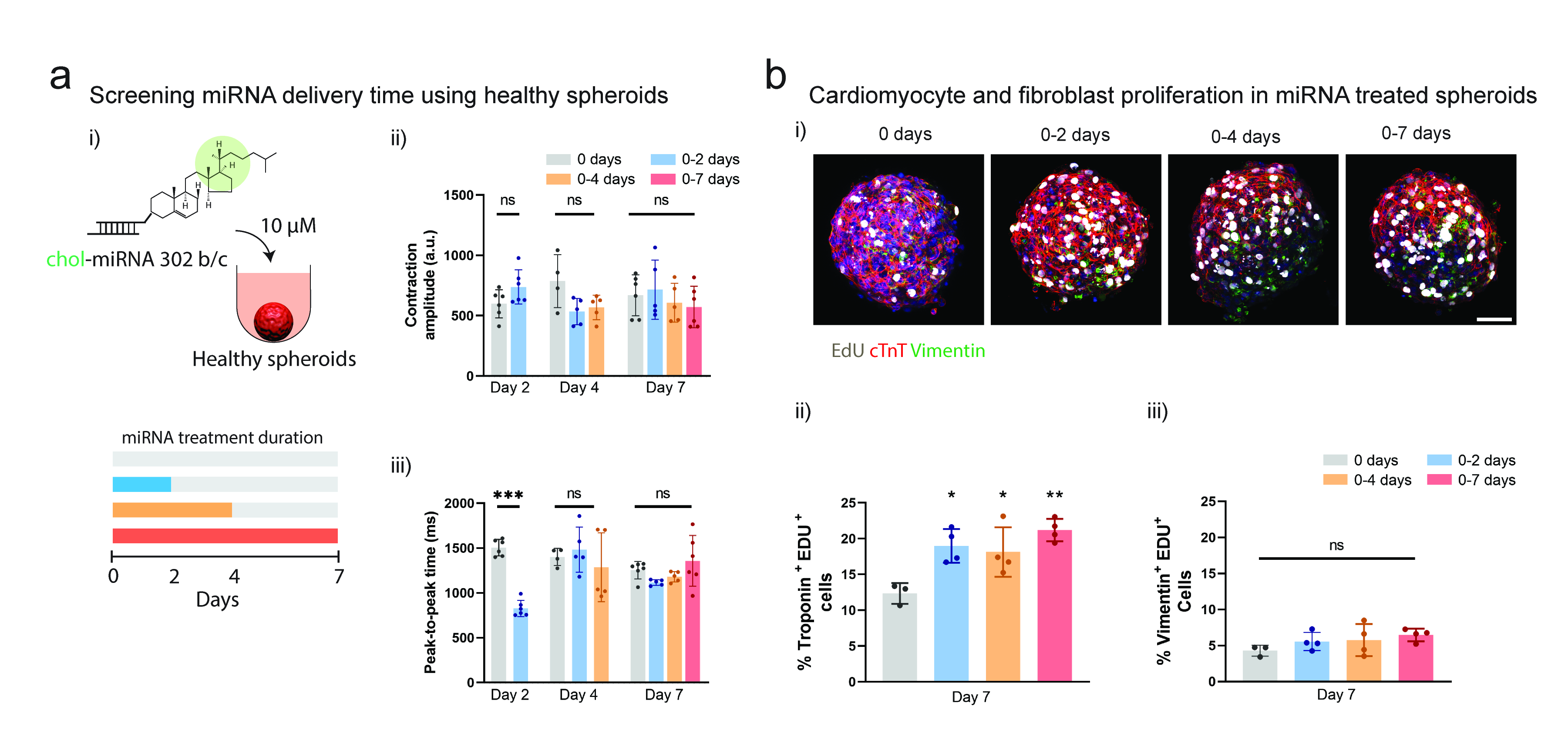

**Supplementary Figure 7: miRNA screening in healthy cardiac spheroids. (a)** (i) Schematic of cholesterol modified miR302 (chol-miRNA 302 b/c) delivery to healthy cardiac spheroids for 0, 0-2, 0-4, and 0-7 days. (ii) Contraction amplitude (a.u) and (iii) peak-to-peak time (ms) within healthy spheroids after 2, 4, and 7 days for each treatment period (n=4-6, mean ± s.d, ns (non-significant), *** p<0.001), one-way ANOVA). **(b)** (i) Immunofluorescence staining for cTnT (red; iPSC-CMs), vimentin (green; cardiac fibroblasts), and EdU (proliferation marker) in healthy spheroids at day 7 for each treatment condition. Quantification of (ii) cardiomyocyte proliferation (EdU^+^ and cTnT^+^) and (iii) fibroblast proliferation (EdU^+^ and Vimentin^+^) at day 7. (n=3-4, mean ± s.d, one-way ANOVA,* p<0.05, ** p<0.01, (0 vs. 0-2 days (p=0.0184): 0 vs. 0-4 days (p=0.0399): 0 vs. 0-7 days (p=0.0026)), ns (non-significant)). Scalebar 50µm.

**Supplementary Movie 1:** Overview of 3D bioprinting process including (i) MSC spheroid aspiration in a media reservoir, (ii) spheroid transfer into a self-healing support hydrogel, and (iii) spheroid deposition within the support hydrogel through removal of vacuum from the capillary tip.

**Supplementary Movie 2:** Fluorescent recording of support hydrogel shear-thinning and self-healing during spheroid transfer through the gel (Note: fluorescent beads (1 µmØ) are embedded in the hydrogel, 1:200 dilution).

**Supplementary Movie 3:** Optical mapping of Ca^2+^ flux in healthy and scarred spheroids after 3 days of culture.

**Supplementary Movie 4:** Brightfield recordings of healthy and scarred microtissue contraction following removal from support hydrogel.

**Supplementary Movie 5:** Optical mapping of Ca^2+^ flux in healthy and scarred microtissues after 5 days of fusion.
